# Quantitative analysis of food web dynamics in a low export ecosystem

**DOI:** 10.1101/2023.03.17.532807

**Authors:** Heather M. McNair, Meredith G. Meyer, Sarah J. Lerch, Amy E. Maas, Brandon M. Stephens, James Fox, Kristen N. Buck, Shannon M. Burns, Ivona Cetinić, Melanie Cohn, Colleen Durkin, Scott Gifford, Weida Gong, Jason R. Graff, Bethany Jenkins, Erin L. Jones, Alyson E. Santoro, Connor H. Shea, Karen Stamieszkin, Deborah K. Steinberg, Adrian Marchetti, Craig A. Carlson, Susanne Menden-Deuer, Mark A. Brzezinski, David A. Siegel, Tatiana A. Rynearson

**Affiliations:** Graduate School of Oceanography, University of Rhode Island, Narragansett, Rhode Island, United States; Department of Earth, Marine and Environmental Sciences, The University of North Carolina at Chapel Hill, Chapel Hill, North Carolina, United States; Department of Cell and Molecular Biology, University of Rhode Island, Kingston, Rhode Island, United States; Institute of Marine Research, Flødevigen, Norway; Bermuda Institute of Ocean Sciences, School of Ocean Futures, Arizona State University, St. George’s, Bermuda; Marine Science Institute, Department of Ecology, Evolution, and Marine Biology, University of California, Santa Barbara, California, United States; Department of Microbiology, Oregon State University, Corvallis, Oregon, United States; College of Marine Science, University of South Florida, Saint Petersburg, Florida, United States; Ocean Ecology Laboratory, NASA Goddard Spaceflight Center, Greenbelt, Maryland, United States; GESTAR II, Morgan State University, Baltimore, Maryland, United States; Monterey Bay Aquarium Research Institute, Moss Landing, California, United States; Department of Botany and Plant Pathology, Oregon State University, Corvallis, Oregon, United States; Department of Oceanography, University of Hawai9i at Mānoa, Honolulu, Hawai9i, United States; Bigelow Laboratory for Ocean Sciences, East Boothbay, Maine, United States; Virginia Institute of Marine Science, William and Mary, Gloucester Point, Virginia, United States; Earth Research Institute & Department of Geography, University of California, Santa Barbara, California, United States

## Abstract

Food webs trace the flow of organic matter and energy among producers and consumers; for pelagic marine food webs, network complexity directly influences the amount and form of carbon exported to the deep ocean via the biological pump. Here we present a synoptic view of mixed layer food web dynamics observed during the late summer 2018 EXport Processes in the Ocean from Remote Sensing (EXPORTS) field campaign in the subarctic Northeast Pacific at the long-running time-series site, Ocean Station Papa. Carbon biomass reservoirs of phytoplankton, microzooplankton, and bacterioplankton, were approximately equal while mesozooplankton biomass was 70% lower. Live organisms composed ∼40% of the total particulate organic carbon within the mixed layer: the remainder was attributed to detritus. Rates of carbon transfer among reservoirs indicated production and assimilation rates were well balanced by losses, leaving little organic carbon available for export. The slight positive net community production rate generated organic carbon that was exported from the system in the form of food web byproducts, such as large fecal pellets generated by mesozooplankton. This characteristically regenerative food web had relatively slow turnover times with small-magnitude transfers of carbon relative to standing stocks that occurred amidst a high background concentration of detrital particles and dissolved organic matter. The concurrent estimation of food web components and rates revealed that separated processes dominated the transfer of carbon within the food web compared to those that contributed to export.

**Plain Language Summary:** The biological carbon pump drives a downward flux of organic matter from the sunlit surface ocean to the vast ocean interior. Ecological interactions in the surface ocean directly affect the amount and type of carbon that is exported to the deep ocean. In this study, we present a synthesis of the late summer mixed layer food web in the Northeast Pacific that was extensively characterized during the 2018 EXport Processes in the Ocean from Remote Sensing (EXPORTS) field campaign. We found the majority of carbon was recycled within the mixed layer by microbes through multiple transfers between producers and consumers. Larger organisms, mesozooplankton and salps, only consumed a small amount of carbon but through the formation of sinking fecal pellets were the main mechanism of transporting carbon out of the system. The study highlights the need to concurrently study microbial and large organism dynamics to develop a predictive understanding of the fate of organic carbon in the oceans.

**Key Points:**

- The microbial loop dominated carbon flow in the late summer mixed layer food web of the North Pacific, most net production was respired leaving little carbon available for export.
- Active production and consumption of organic carbon occurred amid a high background of detrital particulate organic carbon (58% of total) with slow turnover time, 66 d.
- Mesozooplankton which had relatively minor carbon consumption rates created the majority of export production due to efficient repackaging of consumed material.

## 1. Introduction

Carbon flow in oceanic food webs can be characterized by the synthesis of organic carbon by primary producers followed by its consumption and assimilation by a myriad of consumers, and the ultimate conversion of fixed organic carbon to fecal matter, detritus, and respiratory byproducts. Pelagic marine food web processes establish a concentration gradient in organic matter from the sunlit surface to the ocean’s depths that is driven by the downward flux of organic carbon from the surface to the ocean interior, known as the biological carbon pump (BCP) (Michaels and Silver, 1988; Ducklow et al., 2001; Boyd et al., 2019). The complexity of oceanic food webs directly influences both the amount and composition (e.g., phytoplankton cells, zooplankton fecal pellets, dissolved organic carbon (DOC), aggregates) of carbon that contributes to export (Carlson et al., 1994; Durkin et al., 2016; Guidi et al., 2016; McCave, 1975; Passow & Alldredge, 1995; Rynearson et al., 2013; Serra-Pompei et al., 2022; Steinberg & Landry, 2017; Turner, 2015). The links and losses within food webs influence the magnitude and strength of key export pathways within the BCP including sinking of individual phytoplankton cells, sinking aggregates, fecal pellets, and active vertical migration (Nowicki et al., 2022; Siegel et al., 2023). Yet, empirical studies that examine and constrain multiple pathways of export through the food web are rare.

The subarctic North Pacific ecosystem is a model region for understanding low export, regenerative food webs. Extensive time-series programs (e.g., United States and Canadian weather stations and Line P) and large cruise campaigns, (e.g., Canadian JGOFS, SUPER, SERIES) near Ocean Station Papa (Station P) have provided a wealth of information about the long-term variability and seasonal dynamics of this High Nutrient Low Chlorophyll (HNLC) region. The subarctic North Pacific is a relatively physically stable ocean system with modest increases in springtime chlorophyll *a* (chl *a*) concentrations (Philip Boyd & Harrison, 1999; Siegel et al., 2021; Westberry et al., 2016). Primary production is largely fueled by regenerative nitrogen sources (Peña & Varela, 2007; Varela & Harrison, 1999), and production and growth of large cells is limited by iron availability (Boyd et al., 1998; Boyd et al., 1996; Martin & Fitzwater, 1988). Microzooplankton grazing also limits the accumulation of phytoplankton (Boyd et al., 2007; Boyd & Harrison, 1999; Landry et al., 1993; Miller et al., 1991; Strom et al., 1993) though not at all times of year (Rivkin et al., 1999). The biomass of phytoplankton, bacteria, and microzooplankton are often comparable, and fluctuate roughly two-fold over an annual cycle (Booth et al., 1993; Harrison, 2002; Sherry et al., 1999), while the seasonal migration of *Neocalanus* copepods drives a 35-fold change in annual mesozooplankton biomass (Goldblatt et al., 1999). Inverse modeling of the upper ocean food web at this site suggested that irrespective of season, the major trophic pathway of organic carbon within the subarctic North Pacific is from picophytoplankton to microzooplankton to mesozooplankton (Vézina & Savenkoff, 1999).

Only a small fraction of organic carbon production is exported from the euphotic zone from this modestly productive, regenerative food web. Measurements from thorium-234 disequilibrium profiles suggest that particulate organic carbon flux ranges from 3-14% of net primary production (NPP) (Buesseler et al., 2020; Buesseler & Boyd, 2009; Charette et al., 1999) and is primarily composed of fecal pellets with a minor contribution from sinking phytoplankton cells (Durkin et al., 2021; Stamieszkin et al., 2021; Steinberg et al., 2022; Thibault et al., 1999). Comparisons between annual net community production and net particulate flux suggest seasonal contributions to export from DOC and active vertical migration of zooplankton (Bif & Hansell, 2019; Emerson, 2014; Timothy et al., 2013). Collectively, these decades of research provide important insight into the specific environmental and food web components that drive the biological pump in the subarctic North Pacific. Many of these relationships are derived from short-term studies (∼days) made at different times of the year that each address only a subset of food web relationships, requiring inferences relative to carbon export to be derived from the ensemble. To gain quantitative and mechanistic understanding of food web processes and components, concurrent analyses of the major carbon stocks and transfer pathways are needed in combination with their linkages to export flux.

Here we present a synoptic view of the surface ocean ecosystem in the subarctic Northeast Pacific over 28 days in the late summer of 2018. We aimed to 1) characterize the late summer subarctic Northeast Pacific mixed layer food web by quantifying stocks and transformation rates of carbon and compare the dynamics to existing data and models, 2) provide insights into food web variability on both daily and monthly time scales, and 3) quantify the contribution of specific export pathways out of the mixed layer. We find that, consistent with previous studies, short-term oscillations in production and loss are balanced over the month. This leads to a highly retentive food web, characterized by slow turnover times and high levels of carbon recycling and respiratory losses, with no evidence of accumulation of living biomass nor the export of living cells. The primary mechanism that drives carbon export is grazing by mesozooplankton and salps which removes carbon from the recycling microbial loop and repackages small particles into large, sinking fecal pellets.

## 2. Methods

The North Pacific EXPORTS campaign took place from August 15 to September 7, 2018, near Station P (50° N and 145° W). An overview of the geographic scope and physical environment of the campaign is described in Siegel et al. (2021). Here, we synthesize data collected by many EXPORTS researchers to describe the mixed layer food web. The data detail the mixed layer food web observed from the R/V *Roger Revelle,* which sampled in a Lagrangian framework, measuring biological rates and stocks over time. Some of these data have been published in studies focused on specific processes (e.g., Stephens et al., 2020; Maas et al., 2021; Stamieszkin et al., 2021; McNair et al., 2021; Meyer et al., 2022), and all data are available in the SeaBASS or BCO-DMO data repositories (see Table 1 for more detail). Details on each of the methods used to measure food web processes and stocks can be found in the EXPORTS technical memorandum (https://hdl.handle.net/1912/27968, DOI: 10.1575/1912/27968) or in the supplemental information of this manuscript (Table 1).

**Table 1.**
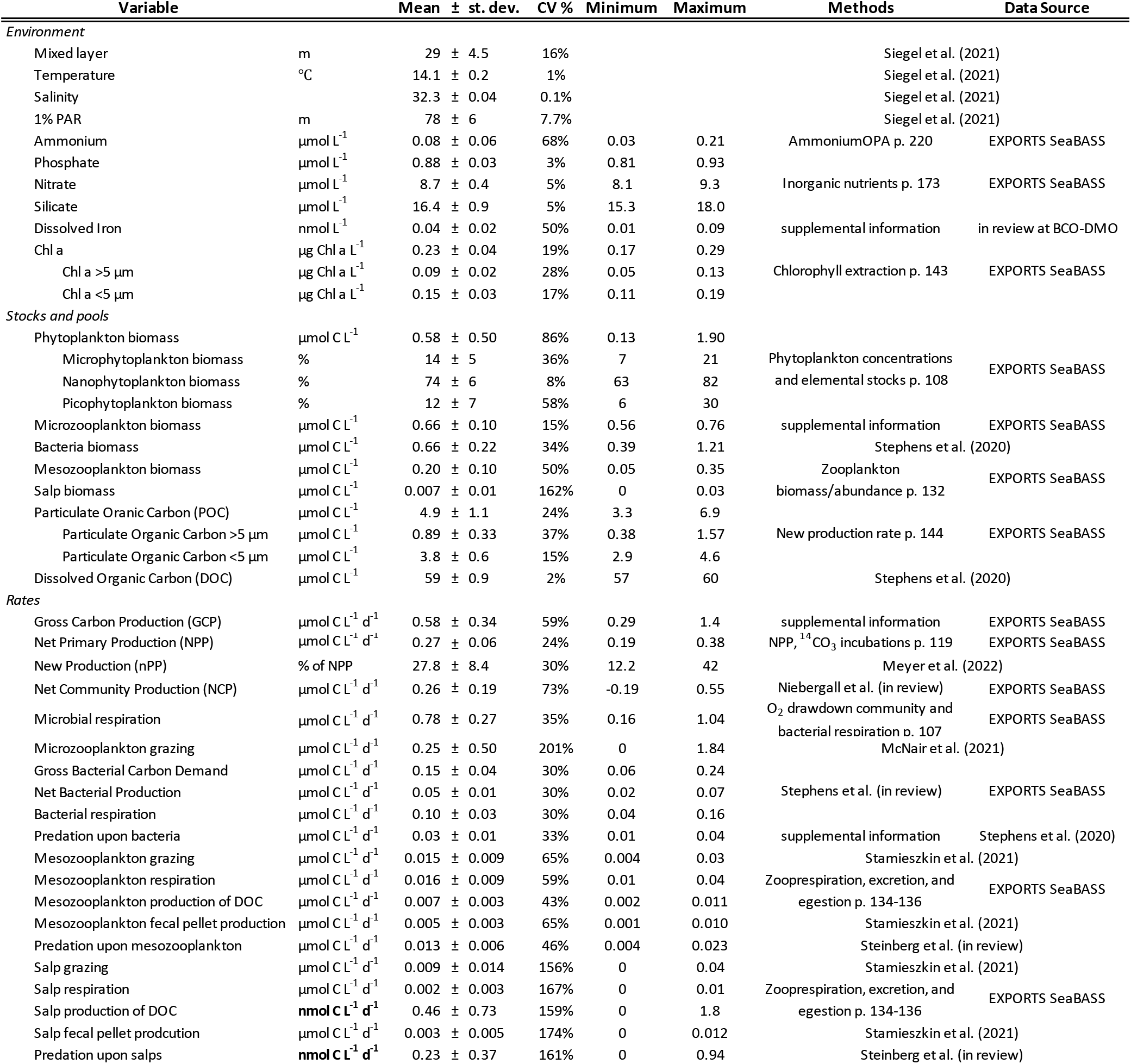
Environmental conditions, and food web stocks and rates. Mean, standard deviation, minimum, and maximum values measured for listed mixed layer food web parameters. Descriptions for the methods of each variable can be found in the citation listed, if the data has been included in a publication, or the parameter name listed in the Methods column. Methods descriptions of all parameter names can be found at https://hdl.handle.net/1912/27968. The primary data from each parameter can be found in the citation listed and in the North Pacific EXPORTS, SeaBass data repository: 10.5067/SeaBASS/EXPORTS/DATA001

The mixed layer depth was defined as the depth where potential temperature was 0.2°C less than the temperature at 5 m (de Boyer Montégut et al., 2004). Data were integrated using trapezoidal integration with values at the mixed layer depth determined using linear interpolation of data that spanned the depth of the mixed layer. Integrated values were then divided by mixed layer depth to remove signal associated solely with changes in integration depth, resulting in weighted averages of mixed layer biomass and rates. All results are presented as mean values with standard deviation unless otherwise noted. While most rate and stock measurements were obtained using just one methodological approach, net community production (NCP) was assessed using nine different methods, across several autonomous and ship-based platforms including on-deck dilution and new production incubation experiments, flowthrough O_2_/Ar, profiling floats and gliders, and via satellite (Niebergall et al., in revision). The average and range of all NCP rate estimates is presented in Table 1.

Primary data sets were converted to carbon units as necessary and carbon assimilation rates were estimated from the literature when no direct measurements were available. Microzooplankton grazing rates on phytoplankton were converted to carbon units using the average mixed layer chl *a* to particulate organic carbon (POC) relationship from the cruise. This conversion was consistent with the balanced growth and grazing rates and the constant phytoplankton and chl *a* stock observed throughout the cruise (McNair et al., 2021) even though it included POC that did not contain chl *a*. Microzooplankton grazing rates on bacterial biomass were estimated from dilution experiments conducted during the cruise (Supplementary Information) and based on data reported in Stephens et al. (2020). Secondary microzooplankton production was not directly measured, therefore a 30% growth efficiency was assumed based on literature estimates of 30-40% (Landry & Calbet, 2004). Size-fractionated biomass of mesozooplankton was calculated using methods described in Steinberg et al. (2008).

Grazing rates of mesozooplankton were calculated from fecal pellet production rates, assuming an assimilation efficiency of 66% (Abe et al., 2013; Steinberg & Landry, 2017). The byproducts from grazing and metabolic activities by the mesozooplankton community and salps were estimated using measured biomass, abundance, and locally validated allometric relationships (Table 1); predation rates upon mesozooplankton were based on allometric estimates of metazoan plankton predator-prey interactions (Zhang & Dam, 1997).

Bacterial carbon production was determined from ^3^H leucine incorporation rates (Stephens et al., in press) using a combination of a previously established ^3^H leucine-to-cell biovolume conversion for Station P (Kirchman, 1992) and a cell biovolume-to-carbon relationship established with samples collected during the cruise (Stephens et al., 2020).

The trophic positions, the number of steps separating an organism from the base of the food web, of micro- and mesozooplankton were calculated by comparing the δ^15^N values of the amino acids alanine and phenylalanine in size-fractionated zooplankton, as in Décima and Landry (2020) and Shea (2021). This alanine-phenylalanine based estimate of trophic position is inclusive of protistan heterotrophy in the underlying food web. The trophic position of protistan heterotrophy was determined by comparing the alanine-phenylalanine based trophic position with glutamic acid-phenylalanine based trophic position (Chikaraishi et al., 2009, exclusive of protistan heterotrophy) of mesozooplankton (Supplementary Information).

Turnover times of stocks were calculated by the dividing the stock by the rate of carbon accumulation or loss from the stock. Additionally, the turnover time of detrital POC was calculated using the turnover times of POC, phytoplankton, bacteria, microzooplankton and the concentration of detritus. The concentration of detrital POC was calculated by subtracting the biomass of phytoplankton, bacteria, and microzooplankton from POC. The turnover time for POC is equivalent to the weighted average of the turnover times of phytoplankton, bacteria, microzooplankton, and detritus, where *w*_i_ is the fractional contribution of phytoplankton (p), bacteria (b), microzooplankton (µz), and detritus (d) to the POC concentration measured from a 1L filtration, and *TO*_i_ is the turnover time for each of the constituents.

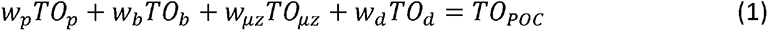

Which can be rearranged to solve for the turnover time of detritus:

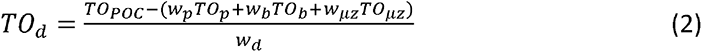

Mesozooplankton are excluded from the detrital measurement because they are not well represented in the 1 L filtered POC measurement. The uncertainly of turnover times was assessed by bootstrapping turnover calculations, using the boot function in R, for 1000 bootstrap cycles and calculating the standard error.

The production of DOC by different components of the food web was determined and compared to bacterial C demand. The amount of DOC produced via phytoplankton extracellular release during gross primary production ranges from 5-35%, with oligotrophic regions tending towards the high end of estimates (Teira et al., 2001). Additionally, 20-40% of the primary production consumed by microzooplankton grazing can be released as DOC (Nagata, 2000; Strom et al., 1997). The fraction of DOC generated by mesozooplankton activities was estimated by applying directly measured allometric relationships (Maas et al., 2021) to the measured abundance and biomass of the mesozooplankton community, while the contributions to DOC from microzooplankton grazing and primary production were estimated from the literature.

To assess correlations among stocks and processes in the food web and to better understand drivers of covariance, a Pearson’s correlation matrix was determined for daily measurements of food web variables with three or more co-occurring data points, i.e., measured on the same day, limiting the dataset to consideration of: macronutrients, DOC, POC, chl *a*, bacterial biomass, new production (nPP), net primary production (NPP), NCP, bacterial production, microzooplankton grazing, and microbial respiration. To visualize relationships between variables, a hierarchical clustering analysis based on Euclidean distances was conducted using the Pearson’s rho value matrix. The clustering analysis was performed with a 12-day subset (every-other day of the cruise) of the data with the highest simultaneous measurement of parameters. Any parameter that was not measured on at least seven of the 12 selected days was not included in the analysis. Data points were extrapolated for the parameters as needed: if data were missing on one of the 12 days, either the nearest neighboring data point was used (only +/- 1 day) or, if two data points were equally spaced, the missing data were extrapolated as the average of the two nearest points. No more than two data points were extrapolated per parameter. The pvclust (Suzuki & Shimodaira, 2006) function in R (R Core Team, 2019) was used to determine statistically significant clusters (p-value <0.05) based on the bootstrap resampling method of Shimodaira (2004). All other statistical analyses and visualizations were conducted in Matlab R2019a using the corrcoef, pdist, linkage (ward method), and dendrogram functions.

## 3. Results

### 3.1 Environmental Overview

Oceanographic and chemical parameters throughout the 2018 field campaign fluctuated around relatively stable cruise-means (Table 1). Atmospheric conditions were generally cloudy, with incident photosynthetically active radiation (PAR) ranging from 10 to 40 moles photons m^-2^ d^-1^ and there were no major storms or other physical perturbations (Siegel et al., 2021). The average euphotic zone (1% PAR) depth was 78 ± 6 m and the mixed layer spanned the upper 40% of the euphotic zone and averaged 29 ± 4.5 m (Siegel et al., 2021). As expected in this HNLC region, the average concentrations of macronutrients in the mixed layer were relatively high, and background dissolved iron concentration generally very low compared to other ocean environments (Table 1).

### 3.2 Food web

The food web was divided into six discrete, biological ‘carbon reservoirs’ (Figure 1). These biological reservoirs, in part, comprise the POC pool (Figure 1). Five of the six carbon reservoirs were quantified: bacteria, phytoplankton, microzooplankton, mesozooplankton (222 µm net), and salps (colored outlined boxes in Figure 1, size of box scales to biomass); the sixth, higher order predators, was not. The flow of carbon between the food web and the POC and DOC pools (colored arrows in Figure 1, width scaled to rate magnitude) is mediated by metabolic and biological rate processes performed by organisms within each carbon reservoir. The waste products of these processes redistribute consumed carbon to the non-living portion of POC (e.g., fecal pellets, carcasses, molts, and aggregations of dead cells), dissolved inorganic carbon (DIC), and DOC (Figure 1). DOC was, by far, the largest pool of organic carbon in the mixed layer–roughly 12 times the size of the POC pool (Table 1); however, ∼95% of the DOC was recalcitrant (Stephens et al., 2020).

**Figure 1.**
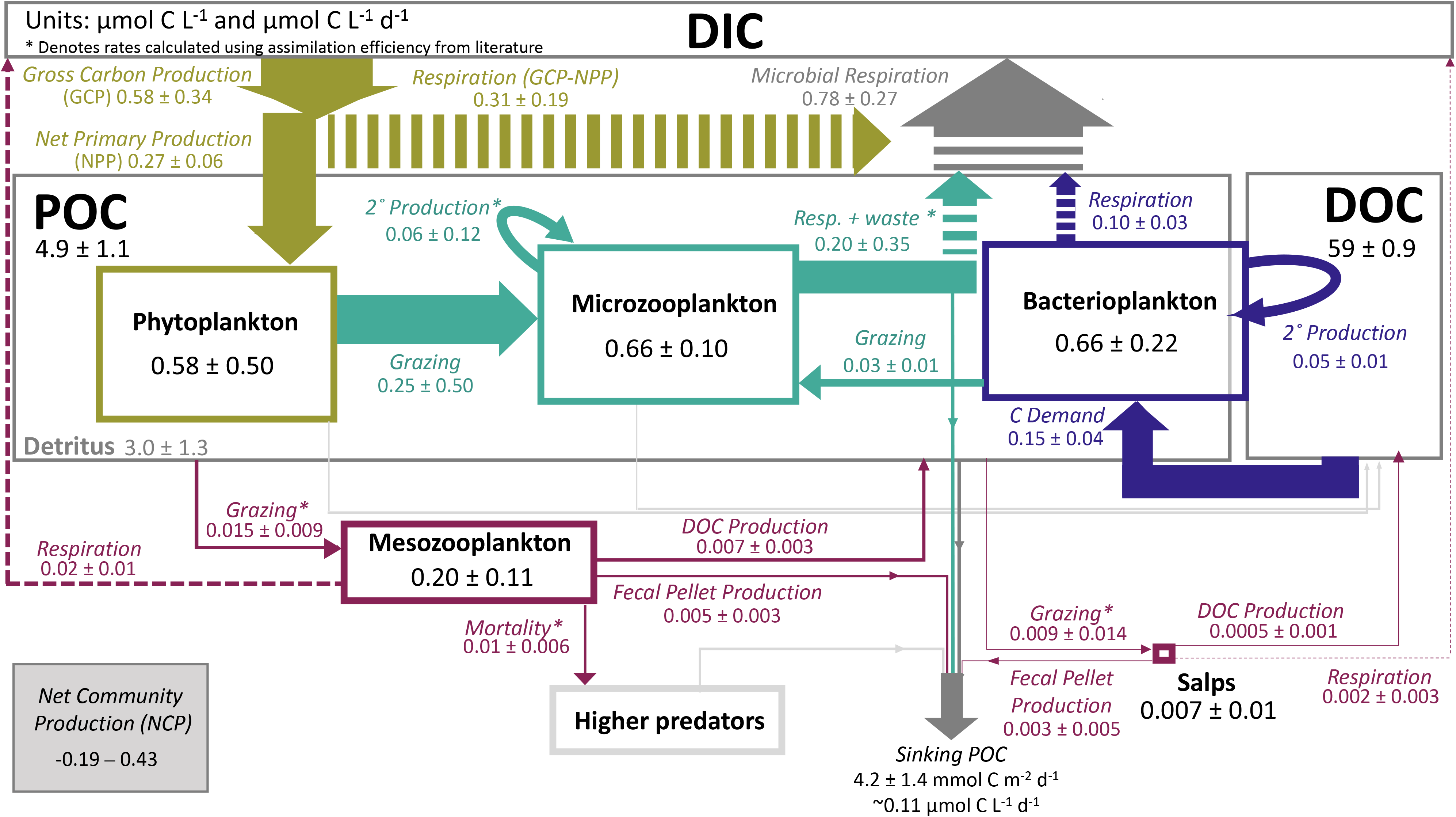
Wiring diagram depicting flows and distribution of carbon within the mixed layer food web. The carbon-based food web is visualized as a set of bulk carbon stocks (DIC, POC, DOC: grey open boxes) and biological carbon stocks (colored, open boxes) connected by rates (filled arrows) of carbon transformation. All values represent cruise averages ± standard deviation and are in units of µmol C L^-1^ (plain text labels) or µmol C L^-1^ d^-1^ (italic labels). Arrow width is scaled to the magnitude of the rate, box area approximates stock concentration, except for DOC, which was 12 times greater than POC. Note the small salp box with adjacent salp label. Stocks and rates are associated with primary producers (green), microzooplankton (teal), mesozooplankton and higher predators (burgundy), and bacteria (blue). Arrows indicate bulk carbon transfers from the POC pool (dark grey), unmeasured contributions to the DOC and POC pools from phytoplankton and microzooplankton stocks via respiration, exudation, and other processes (thin, pale grey) and carbon transfers into the DIC pool, i.e., respiration (dashed). Some rates were calculated using assimilation efficiencies from literature (asterisks). Phytoplankton, microzooplankton and bacteria stocks are subsets of the total POC stock. The concentration of detrital POC was calculated as total POC minus the biomass of phytoplankton, microzooplankton, and bacteria.

#### 3.2.1 Biomass distribution in the food web

The average mixed layer concentration of POC, measured from 1 L filtration volumes, during the cruise was 4.9 ± 1.1 µmol C L^-1^, 84% of which was <5 µm in diameter. Adding the mesozooplankton and salp biomass from MOCNESS tows, which is not accurately represented in a 1 L filtration, brings the total POC concentration to 5.1 ± 1.1 µmol C L^-1^ (Figure 1). Independent measures of organismal biomass indicate that the living fraction of POC was fairly evenly divided among primary producers (11%), consumers (17% total: 13% microzooplankton, 4% mesozooplankton) and recyclers (bacterioplankton, 13%) (Figure 2). Organismal biomass only comprised 41% of the POC pool, presumably the remaining 59% was detrital POC. However, this may be an underestimate of the detrital POC pool given that ∼50% of bacterioplankton cells pass through the precombusted GF/F filters used to sample bulk POC (Lee et al., 1995) and that our accounting of organismal biomass POC includes all bacterial biomass.

**Figure 2.**
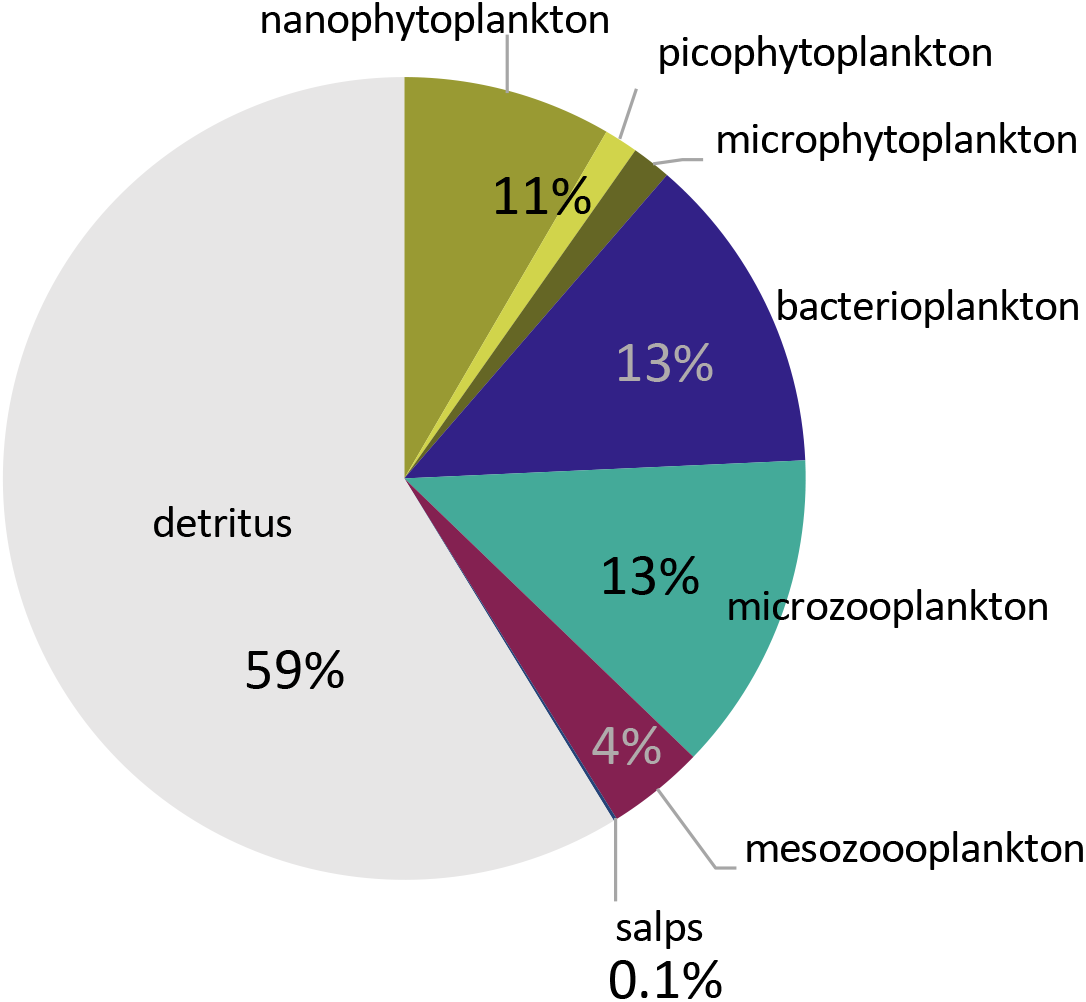
Distribution of particulate organic carbon. The percent contribution of identified particulate organic carbon (POC + mesozooplankton biomass) within the mixed layer food web. The uncharacterized portion of POC (59%, gray area) is assumed to be detrital.

The phytoplankton community was dominated by small cells <5 µm in diameter that made up 65% of the chl *a* stock, including the numerically dominant cyanobacterium *Synechococcus* (McNair et al., 2021; Sharpe et al., 2022). Small cells also dominated phytoplankton carbon stock: 74 ± 6% of the biomass was composed of nanophytoplankton, 12 ± 7% was picophytoplankton, and 14 ± 7% was microphytoplankton (Table 1 & Figure 2) as determined via flow cytometry. The eukaryotic phytoplankton community was primarily composed of species from the genera *Phaeocystis, Pseudochattonella, Chrysochromulina*, *Pseudo-nitzschia, Aureococcus,* and *Plagioselmis,* determined using amplicon sequencing.

The carbon biomass of the microzooplankton community, as determined using cells sizes from microscopy, was composed of ciliates (63%) and dinoflagellates (37%). Taxa from the order Strombidiida, and dinoflagellate genera, *Karlodinium* spp. and *Gymnodinium* spp were abundance in the 18S rDNA amplicon sequencing data.

The mesozooplankton community was dominated by crustacean zooplankton (such as *Neocalanus* spp. copepods) and sporadically by a salp bloom (*Salpa aspera;* Steinberg et al. 2022). The diel migratory community, present in the mixed layer only during the night, consisted primarily of calanoid copepods including *Metridia pacifica,* salps, and the ontogenetic migrators *Neocalanus cristatus* and *N. plumchrus*. Although abundant members of the community, the ontogenetic migrators were entering diapause during the cruise, resulting in very low fecal pellet production and inferred grazing rates by these organisms (Stamieszkin et al., 2021). Other dominant migrators included amphipods, particularly *Themisto pacifica* and *Vibilia propinqua,* as well as a diverse assemblage of euphausiids, several large migratory chaetognaths, and a few large *Clio pyramidata* thecosome pteropods.

For bacterioplankton, both metagenome and 16S rDNA amplicon sequencing identified members of the *Alphaproteobacteria*, (particularly SAR11 and Roseobacter clade), *Gammaproteobacteria*, and *Bacteroidetes* classes to be abundant in similar proportions of 20- 30% per class (Stephens et al., in press; Sharpe et al., personal communication). Cruise-based experiments also found significant increases in the relative abundance of a diverse array of bacterioplankton taxa including members of the *Methylophilaceae* family (OM43 genus) and KI89A order, as well as members of *Bacteroidetes* (*Flavobacteriaceae* NS2b genus), *Alphaproteobacteria* (*Rhodobacteraceae: Sulfitobacter* genus), and *Gammaproteobacteria* (*Alteromonadales order* and *Ectothiorhodospiraceae family*) classes (Stephens et al., 2020). *3.2.2 Biological rates* Net primary production was on average 47% of gross carbon production (GCP). An average of 28% of NPP was nPP, defined as primary production supported by nitrate uptake (Meyer et al., 2022). Microzooplankton grazing on phytoplankton balanced the rate of NPP when averaged over the cruise (McNair et al., 2021). In contrast, the microzooplankton grazing rate on bacteria was roughly 60% of the net bacterial production rate (Table 1). Results of isotope analysis of individual amino acids indicated that the trophic position of heterotrophic protists spanned between the second and third levels in the food web, with the number of trophic steps within the group being 1.4 ± 0.8 (Shea, 2021). Thus, net microzooplankton production was estimated to be 0.06 ± 0.12 µmol C L^-1^ d^-1^ given 1.4 trophic steps and a 30% growth efficiency (Landry & Calbet, 2004). The average rate of bacterioplankton production was greater than the combined losses due to respiration and grazing, leading to an implied accumulation rate of 0.02 ± 0.01 µmol C L^-1^ d^-1^ of biomass (Figure 3c).

**Figure 3.**
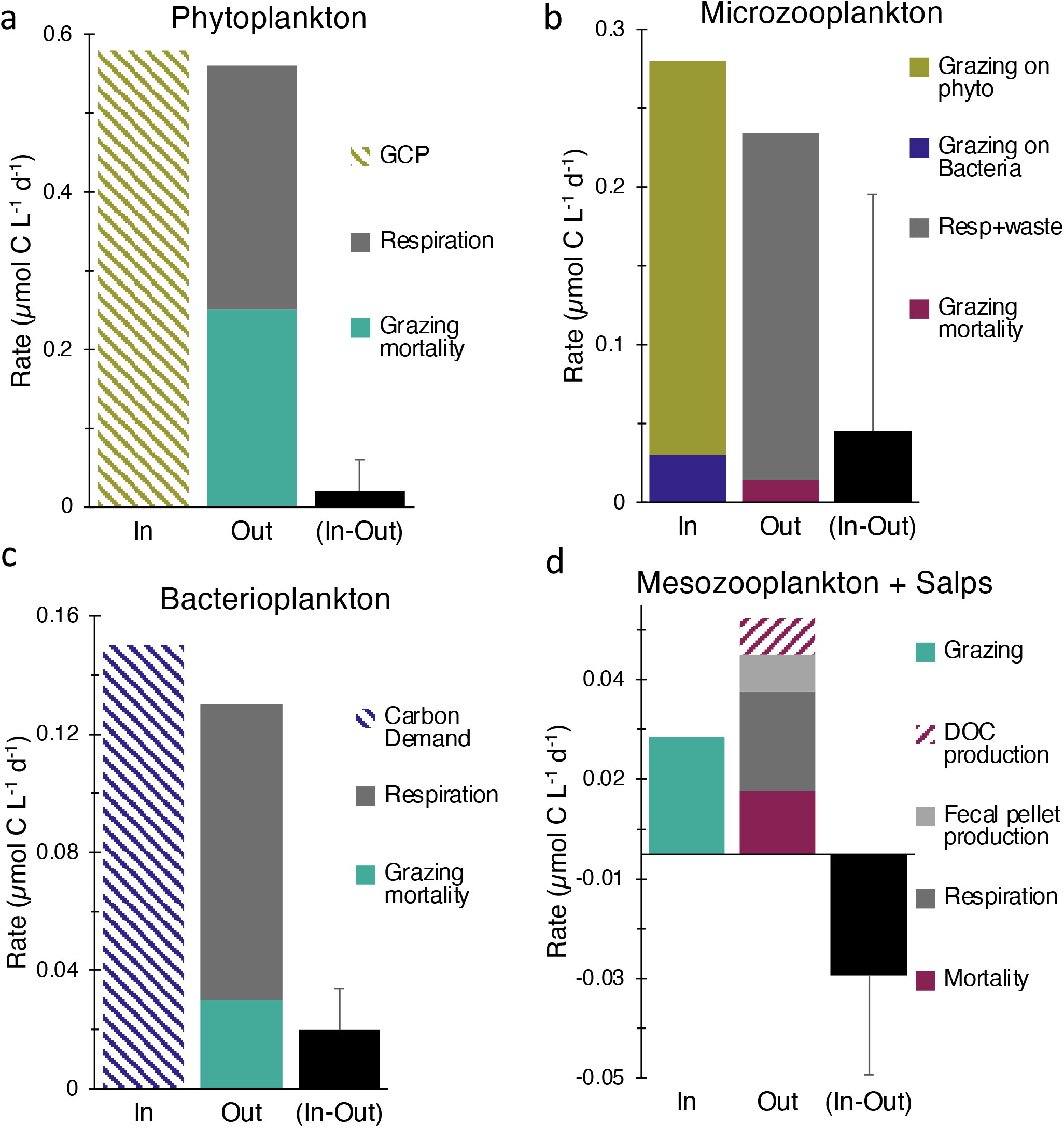
Balance of carbon flows through biological components of the food web. Colored and textured stacked bars reflect the measured, and estimated, mean rates of carbon uptake and production (In) versus carbon consumption, loss and mortality (Out) for the biological components of the food web: phytoplankton (a), microzooplankton (b), bacterioplankton (c), and mesozooplankton + salps (d). The net difference between the combined In and Out rates for each biological group are shown as black bars with error bars representing propagated standard deviation and include biological and analytical variability. Abbreviations are as follows: gross carbon production (GCP), respiration (resp), dissolved organic carbon (DOC).

Feeding by mesozooplankton on primarily microzooplankton and detritus was relatively low, roughly 5% of the rate of microzooplankton grazing (Table 1). Mesozooplankton metabolic byproducts were distributed among particulate and dissolved carbon pools, with 61% to DIC, 16% to POC and 23% to DOC (Figure 1, Table 1). Roughly 6% of the mesozooplankton community biomass was lost daily to predation by larger metazoans. Salp metabolic byproducts (DOC and respiration) were an order of magnitude less than mesozooplankton. Yet, salp fecal pellet production, was similar to that of mesozooplankton (Maas et al., 2021; Stamieszkin et al., 2021; Steinberg et al., 2022).

Mixed layer bacterial growth efficiencies were 31% on average (Stephens et al., 2020). These efficiencies were combined with ^3^H-Leucine incorporation-based estimates of net bacterial production to estimate a cruise mean bacterial carbon demand of 0.15 µmol C L^-1^ d^-1^. Based on bacterioplankton incubations, most of the DOC was recalcitrant on the time scale of weeks, with only ∼5% (3 µmol C L^-1^) of the DOC consumed by bacterioplankton (i.e., “bioavailable” fraction) over ∼90 days (Stephens et al., 2020).

### 3.3 Balance between production and loss

The relative balance of carbon uptake and loss by each food web reservoirs provides a map of the biological processes, and their magnitudes, that potentially contribute to carbon export. The concentration of carbon did not significantly change over the course of the cruise in any of the particulate reservoirs (linear regression, all p-values > 0.15). Only 1-21% of the variability in the POC reservoirs was linearly dependent on time (R^2^ = 0.01 to 0.21). It is thus expected that the rates of production and loss were balanced for each reservoir over the timeframe of the cruise. Phytoplankton production and losses were well balanced over the course of the cruise (Figure 3a). GCP was balanced by phytoplankton respiration (GCP-NPP), and grazing (McNair et al., 2021), resulting in no statistically significant net accumulation or loss of phytoplankton biomass (linear regression R^2^= 0.02, p-value = 0.15). Grazing upon phytoplankton was overwhelmingly dominated by microzooplankton whose grazing rate on phytoplankton exceeded the grazing rate of mesozooplankton on all forms of carbon by a factor of 16 (0.25 µmol C L^-1^ d^-1^ ÷ 0.015 µmol C L^-1^ d^-1^).

Although not quantified directly, consumption of microzooplankton biomass is carried out by higher trophic level zooplankton like salps, euphausiids, and non-ontogenetically migrating copepods (Landry & Calbet, 2004) and thus is included in the mesozooplankton grazing rates. Some inference as to the place of secondary consumers within the food web was gained through amino acid isotope analysis that placed mesozooplankton in trophic positions 3.5 and 4.5 within the food web with larger mesozooplankton (>5 mm) occupying the higher trophic position (Shea, 2021). The estimated net production of microzooplankton biomass is roughly three-fold higher than the carbon requirements of mesozooplankton, but variability in razing and predation rates create an estimate of net growth with error bars that span zero (Figure 3b).

Mesozooplankton rates of organic carbon gain and loss were unbalanced and suggest a decrease in mesozooplankton biomass during the study period. Mesozooplankton losses (respiration, DOC production, POC production, and predation mortality) were roughly three-fold higher than the amount of carbon consumed by mesozooplankton, leaving a net removal rate of 0.03 µmol C L^-1^ d^-1^ of mesozooplankton biomass (Figure 3d). Salp presence was highly variable (coefficient of variation, coefficient of variation = 162%, Table 1), but overall, the rate of carbon consumption and loss for salps was balanced.

The diversity of methodological approaches employed during EXPORTS allowed us to estimate DOC production by various components of the food web and compare this to bacterial carbon demand (i.e., gross bacterial production). We estimated DOC release rates of 0.03-0.20 µmol C L^-1^ d^-1^ by phytoplankton, 0.05-0.10 µmol C L^-1^ d^-1^ by microzooplankton, and 0.007 ± 0.003 µmol C L^-1^ d^-1^ by mesozooplankton during the cruise. These estimates were summed to generate an estimated 0.08 - 0.30 µmol C L^-1^ d^-1^ of total DOC production which encompasses the measured bacterial carbon demand of 0.15 ± 0.04 µmol C L^-1^ d^-1^.

In addition to examining the balances of the reservoirs individually, we analyzed the carbon balance of the total mixed layer food web (Figure 4a). We compared the balance between community respiration (CR) to GCP using two approaches. First, we determined O_2_ drawdown in seawater to represent the combined respiration of phytoplankton, bacteria, and microzooplankton. It averaged 0.78 ± 0.27 µmol C L^-1^ d^-1^ and when combined with respiration from salps and mesozooplankton, the community respiration rate was 0.8 µmol C L^-1^ d^-1^ (Figure 4a, CR 1). We obtained a second estimation of CR by summing the measured respiration of phytoplankton (i.e., the difference between GCP and NPP), bacteria, and mesozooplankton, and adding an estimated microzooplankton respiration rate of ∼50% of consumed carbon (Calbet & Landry, 2004) yielding a community respiration rate estimate of 0.57 µmol C L^-1^ d^-1^ (Figure 4a, CR 2). The second estimate of community respiration is ∼30% lower than the first but within the variability of the directly measured rate of microbial respiration (0.16-1.14 µmol C L^-1^ d^-1^). The carbon lost from the food web by respiration is thus approximately equivalent to the carbon entering the system as GCP whether compared to the sum of group-specific respiration rates (0.56 µmol C L^-1^ d^-1^) or the sum of the directly measured microbial, mesozooplankton, and salp respiration rates (0.80 µmol C L^-1^ d^-1^), with GCP:CR (community respiration) ranging from 0.73 to 1.02.

**Figure 4.**
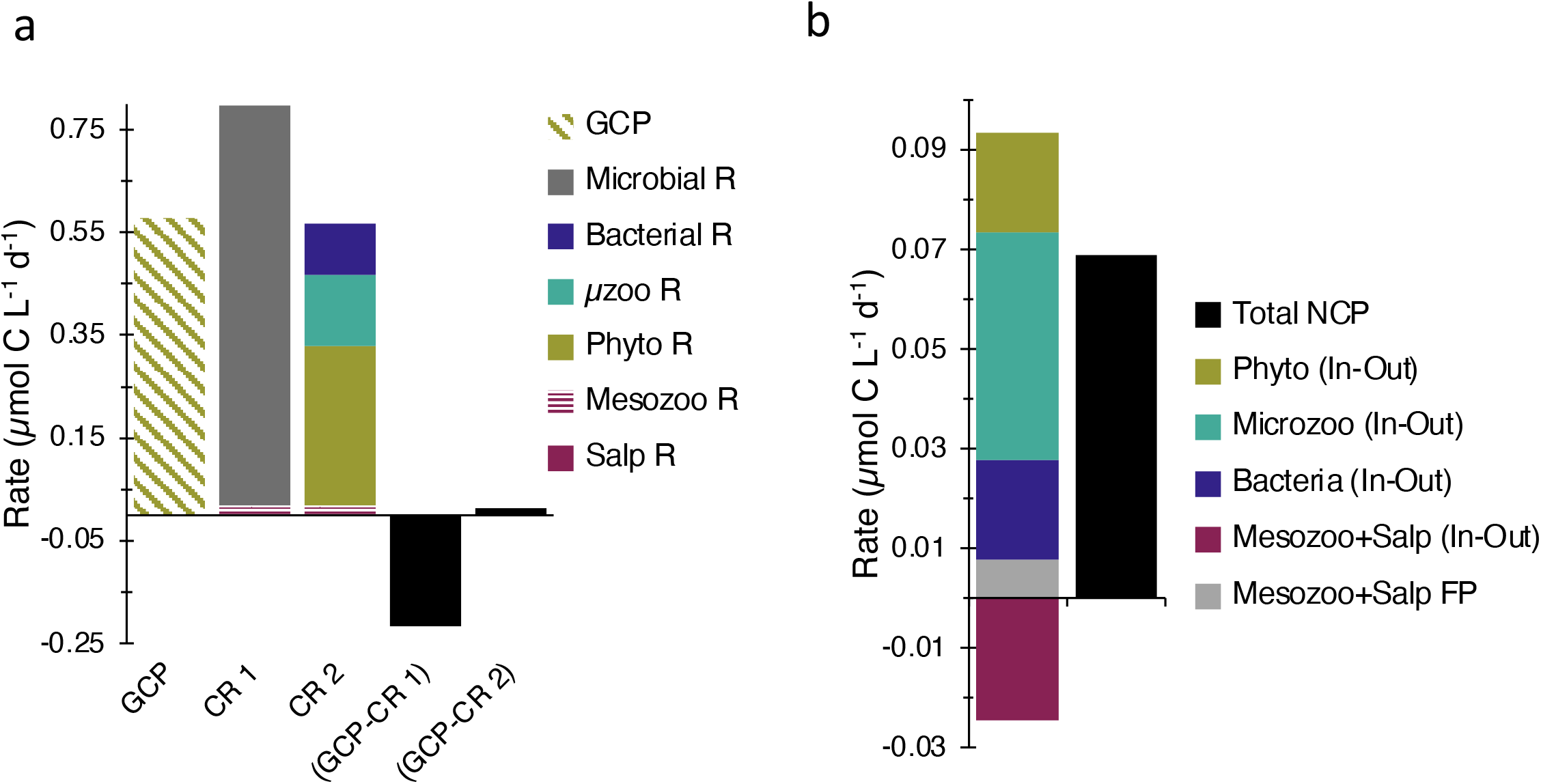
Balance between gross carbon production and community respiration. (a) Phytoplankton gross carbon production (GCP) compared with two estimates of community respiration. The first estimate of community respiration (CR 1) combines microbial respiration (i.e., the respiration rate measurement from unfiltered seawater which includes phytoplankton, microzooplankton and bacteria), mesozooplankton (Mesozoo R), and salp respiration (Salp R). The second estimate of community respiration (CR 2) combines the respiration rate of phytoplankton (Phyto R), microzooplankton (µzoo R), bacteria (Bacterial R), mesozooplankton and salps. The black bars show the difference between GCP and CR 1, -0.22 ± 0.38 µmol C L^-1^ d^-1^, and the difference between GCP and CR 2, 0.01 ± 0.13 µmol C L^-1^ d^-1^. (b) Net community production (NCP, 0.07 ± 0.27 µmol C L^-1^ d^-^1, black bar) is the sum of the In-Out for each of the biological components from Figure 3 in addition to the contribution of mesozooplankton and salp fecal pellets (Mesozoo+Salp FP). Note the y-axes are scaled differently for a and b.

While the production and loss of organic carbon from the food web were equivalent within measurement error, results from thorium-234 profiles and sediment traps indicate a net production of carbon that contributes to flux out of the mixed layer (Buesseler et al., 2020; Durkin et al., 2021; Estapa et al., 2021; Roca-Martí et al., 2021). The sum of the difference between the production and loss of each biological carbon reservoir (In-Out, Figure 3) plus the contribution of fecal pellets (fp) from mesozooplankton and salps yields an estimate of net community production of 0.07 ± 0.27 µmol C L^-1^ d^-1^ (Figure 4b) with a standard deviation propagated from the individual measurements. Relationships between fecal pellet production and grazing rate are not available for microzooplankton; however, the estimate of net production would increase by roughly 10% if 30% of microzooplankton-grazed carbon was converted into fecal pellets. The above estimate of net community production from individual food web measurements is roughly 1/4^th^ of the average NCP of 0.26 µmol C L^-1^ d^-1^ measured independently among multiple autonomous sampling platforms, e.g., gliders, floats, incubations etc. (see Siegel et al., 2021), but falls within the range of estimates provided by these platforms (-0.19–0.55 µmol C L^-1^ d^-1^) (Niebergall et al., in revision). Integrating the mixed layer NCP rate from the food web analysis (0.07 ± 0.27 µmol C L^-1^ d^-1^) to 40 m yields 2.8 mmol C m^-2^ d^-1^, comparable to the thorium-derived estimate of carbon flux at 40 m, 4.2 ± 1.4 mmol C m^-2^ d^-1^ (Buesseler et al., 2020). Overall, carbon production and consumption in the food web were reasonably balanced and suggest a positive net production on the order of 0.07 µmol C L^-1^ d^-1^ and a maximum of 0.55 µmol C L^-1^ d^-1^ of food web byproducts that could contribute to export.

### 3.4 Turnover times

While we do not assume that all components of the food web were in steady state, we estimated turnover times to examine relative rates of food web mediated carbon cycling during the cruise. Turnover times are presented with bootstrapped standard error. Phytoplankton stocks turned over every 2.1 ± 0.57 d (Figure 5). Turnover time of microzooplankton stocks was five-fold longer, 11 ± 16 d and similar to the turnover time of bacteria in the mixed layer, 14 ± 0.65 d. The turnover time of POC in the mixed layer was estimated using the ^234^Thorium- derived particle flux at ∼40 m (4.2 ± 1.2 mmol C m^-2^ d^-1^) (Buesseler et al., 2020) and the POC content in the upper 40 m (Table 1), yielding a turnover time of total POC in the mixed layer of 44 ± 3.1 d. The turnover time of the detrital POC was 66 ± 6.3 d. The turnover time of the bioavailable fraction of DOC was 20 ± 5.3 d and represents the actively used portion of the DOC pool measured experimentally during the cruise (Stephens et al., 2020). Turnover times for the recalcitrant portions of the DOC pool are on timescales of years to millennia (Carlson & Hansell, 2015). Turnover time was not estimated for mesozooplankton because it is less informative for higher trophic level organisms with multiple life stages and more complex life histories that span timescales longer than the cruise occupation.

**Figure 5:**
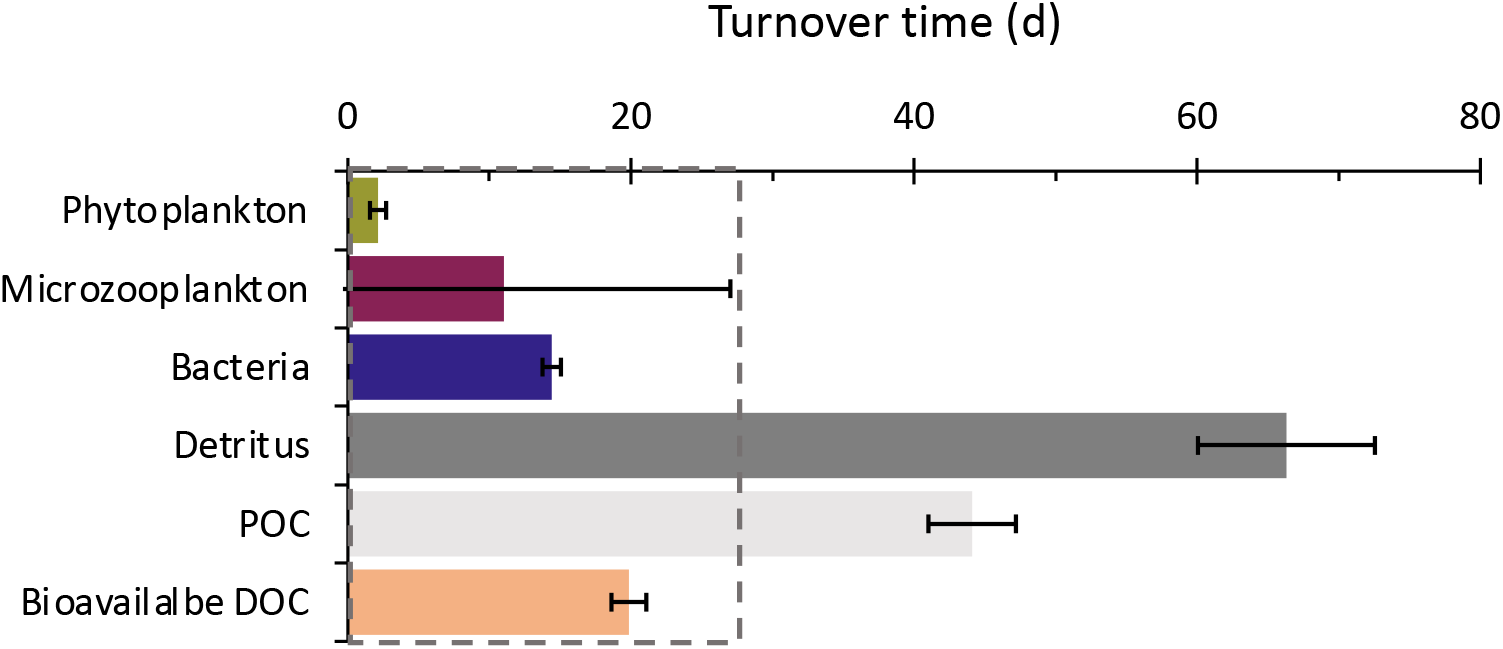
Turnover times in the mixed layer food web. The turnover times in days (d) of organic carbon stocks in the mixed layer food web. Dashed box shows the duration of the cruise. Abbreviations are as follows: particulate organic carbon (POC), dissolved organic carbon (DOC). Error bars show the standard error of 1000 bootstrap cycles.

### 3.5 Production and loss rates of detritus

The extensive measurements of stocks and rates enabled us to infer some of the dynamics of detritus in the food web, including micro- and mesozooplankton fecal pellets, dead cells, and fragments of aggregates and larger particles. Detrital POC was 59% of the total POC in the mixed layer, yet its long turnover time of 66 d yields a relatively slow rate of detrital production, consumption, and loss of ∼0.05 µmol C L^-1^ d^-1^ (3 µmol C L^-1^ 66 d). While mesozooplankton biomass is not well represented in 1 L POC measurements, mesozooplankton fecal pellets might be, so we subtracted meso- and salp fecal pellet production rates from the detrital production rate to estimate production of ‘other’ detritus.

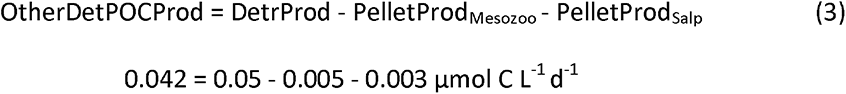

The production rate of ‘other’ detritus is equivalent to the loss of living POC to the detrital pool. To determine if the production rate of ‘other’ detritus could be supported by the living organic carbon pool, we calculated the production rate of living POC (biomass) by accounting for gains and losses of bacteria, phytoplankton, and microzooplankton and obtained a positive production rate of 0.09 µmol C L^-1^ d^-1^.

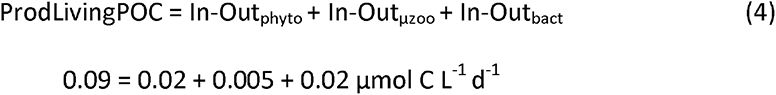

Thus, to support the production of detritus, it seems that most of the ‘net’ living biomass that is created becomes detritus.

### 3.6 Correlations of food web dynamics within the mixed layer

The results presented thus far have focused on cruise-wide averages to establish a mean ecosystem state; the daily variation within these averages provides insight into the scale of coherence between food web parameters (Supplemental Figure 1a). Several food-web variables were measured both simultaneously and sufficiently frequently (Supplemental Figure 1b) to examine their relationships using hierarchical clustering (see methods). This analysis generated four distinctly clustered branches (Figure 6). Each branch represents a group of positively correlated food web variables that similarly fluctuate in relation to the rest of the parameters. Total DOC was not significantly correlated, nor did it show similar correlations patterns with any other food web variables and thus remained distinct and isolated from the other clusters (Figure 6). Cluster A was comprised of POC, chl *a,* and NPP, which are significantly positively correlated (Pearson’s, p value <0.05) to one another and negatively correlated to macronutrient concentration. The next branch (cluster B) shows a cascading cluster of positively correlated variables: NCP, bacterial biomass, net bacterial production, and nPP. Cluster B variables are characterized by relatively weaker positive correlations to cluster A variables and negative correlations to macronutrient concentrations (Supplemental Figure 1b). Microbial community respiration, which contributes to community respiration, and microzooplankton grazing showed similar correlation patterns to other food web variables and composed cluster C. Microzooplankton grazing was significantly correlated to NCP, however the weak positive correlation with macronutrients and negative correlation with NPP kept microzooplankton grazing from being clustered with NCP. Macronutrient concentrations were clustered on a branch distinctly separate from the other branches and were negatively correlated to almost all other food web parameters (cluster D).

**Figure 6:**
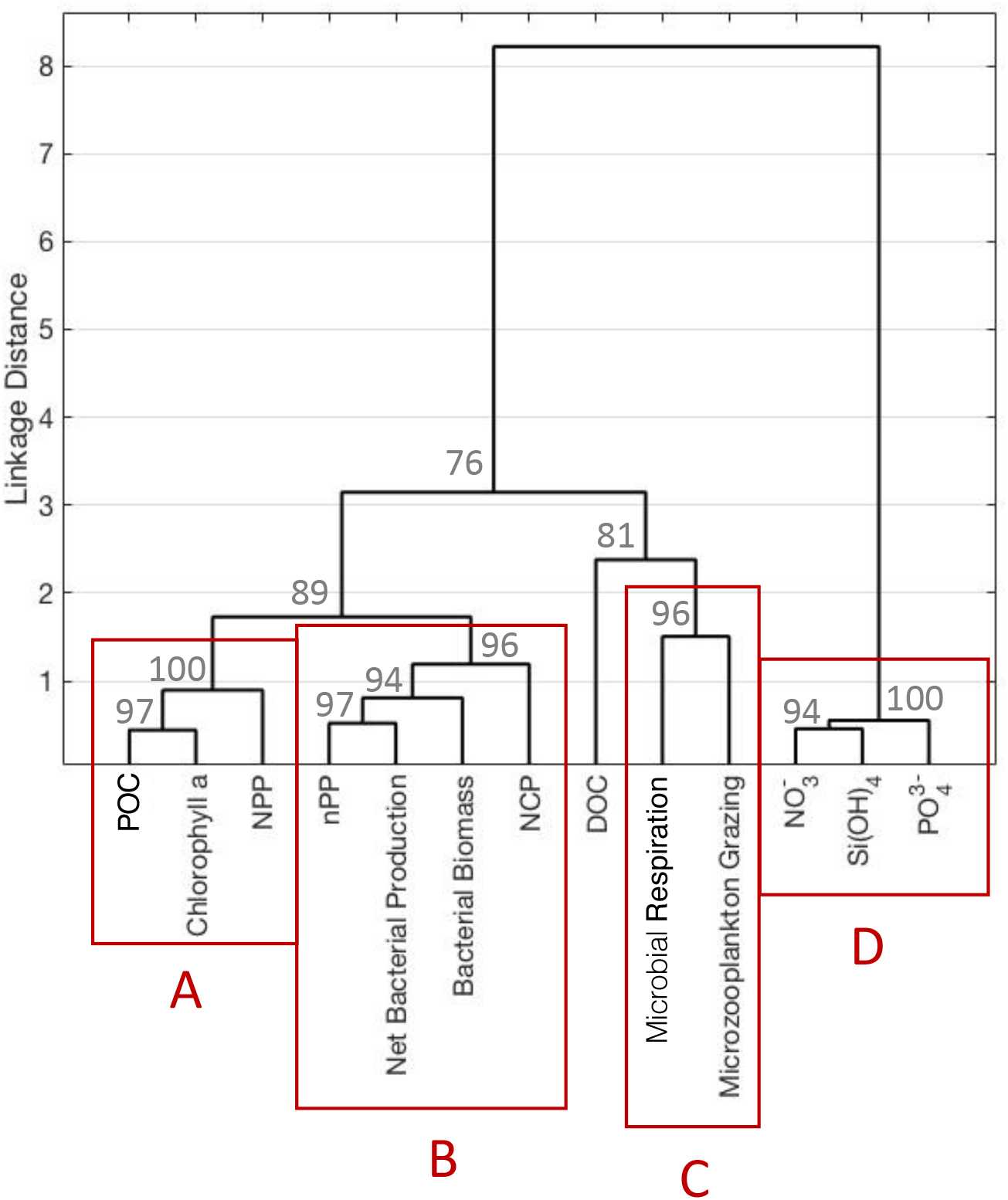
Clustering of mixed layer food web variables. Dendrogram visualizing the linkage distance of food web stocks and rates based on a pair-wise Pearson’s correlation matrix (Supplemental Figure 1b). Confidence levels (1 – p-value) for the clusters (gray text) are at branch nodes. Red boxes show the highest order clusters with significant grouping (A-D, p-value < 0.05). Abbreviations are as follows: net primary production (NPP), chlorophyll a (chl *a*), new primary production (nPP), net community production (NCP), microzooplankton grazing rate (µzoop grazing), integrated daily radiation (I_g_), ortho phosphate (PO ^3-^), microbial community respiration (CR), particulate organic carbon (POC), nitrate (NO ^-^), silicate (Si(OH)_4_), and dissolved organic carbon (DOC).

## 4. Discussion

During the North Pacific EXPORTS campaign, production and consumption were balanced on average which resulted in a highly retentive food web characterized by high levels of carbon recycling and respiratory losses. These dynamics led to relatively slow rates of carbon transfer within the food web that occurred against a large background concentration of persistent dissolved and particulate organic carbon. Carbon that was exported from the system took the form of food web byproducts, such as fecal pellets and detritus (Durkin et al., 2021). Export was primarily facilitated by the sporadic influence of zooplankton whose grazing removed carbon from the recycling microbial loop and repackaged small particles into larger, gravitationally sinking particles (Maas et al., 2021; Stamieszkin et al., 2021; Steinberg et al., 2022).

### 4.1 Ecosystem state

The ecosystem state observed during the August 2018 EXPORTS campaign falls towards the lower range of previously measured carbon production rates and stocks for the region. The physical environment during late summer was consistent with climatological records while biological production rates and standing biomass were generally lower than average and euphotic zone depths were considerably deeper (Siegel et al., 2021). The concentration of POC, DOC, and mesozooplankton biomass were low and closer to historic wintertime conditions (Bif & Hansell, 2019; Goldblatt et al., 1999; Harrison, 2002). The average mixed layer chl *a* concentration was roughly half of the summertime average (Philip Boyd & Harrison, 1999; Siegel et al., 2021), and phytoplankton carbon was roughly one-third of prior late summer estimates (Paul J. Harrison, 2002). Microzooplankton biomass was similar to the lowest measured during the SUPER cruises (Booth et al., 1993), and bacterial biomass was comparable to the low concentrations previously observed during the spring (Kirchman et al., 1993; Sherry et al., 1999).

While carbon stocks were relatively low, the distribution of carbon among most pools was generally consistent with previous late summer observations. The majority of POC was likely detrital and the remainder was distributed evenly among phytoplankton, microzooplankton, and bacterioplankton, as seen previously (Booth et al., 1993; Paul J. Harrison, 2002; Sherry et al., 1999). Relative to the microbial biomass, mesozooplankton biomass was lower than previous late summer observations (Goldblatt et al., 1999; Paul J. Harrison, 2002). The ratio of phytoplankton carbon to mesozooplankton biomass was 3:1, which was greater than the 1:1 ratio seen in late summer but not as high as the ratio of ∼5 observed during winter (Goldblatt et al., 1999; Paul J. Harrison, 2002). DOC concentration was an order of magnitude greater than any other organic carbon pool; however, the majority of DOC was recalcitrant with only a small percentage being bioavailable on time scales of days to weeks (Stephens et al., 2020), supporting previous observations (Carlson 2002, Carlson and Hansell, 2015).

The rates of carbon flow between stocks also fell towards the lower end of previous measurements, but the balanced microbial dynamics were consistent with HNLC food web paradigms (Boyd et al., 2004; Boyd & Harrison, 1999; Miller et al., 1991). A balance between production and loss, and no significant change in the concentration of carbon reservoirs over time suggests the system approximated steady state when processes were averaged over the 28-day cruise, except for declining mesozooplankton biomass. Primary production and bacterial production were within the lower range of previous observations for Station P (Giesbrecht et al., 2012; Kirchman et al., 1993; Marchetti et al., 2006; Miller et al., 1991; Sherry et al., 1999). Consistent with current paradigms for the subarctic Pacific Ocean HNLC region, primary production was mainly fueled by regenerated nitrogen (Meyer et al., 2022), and grazing by microzooplankton limited the accumulation of the dominant picophytoplankton in the mixed layer by consuming them at the same rate they were being produced (McNair et al., 2021). The growth of larger phytoplankton, primarily diatoms, was limited by the availability of Fe (Jenkins, personal communication), consistent with previous iron enrichment experiments (Martin and Fitzwater 1988; Boyd et al. 1996, 1998; Marchetti et al. 2006).

Mesozooplankton biomass was in a period of decline with respiration and excretion rates that exceeded ingestion rates. The region is well known to have a seasonal pattern of mesozooplankton biomass, with a peak during the spring bloom and a decline to a winter low (Goldblatt et al., 1999; Mackas & Galbraith, 2002). The mesozooplankton grazing rate and fecal pellet production rates were lower than previous summer measurements (see Stamieszkin et al., 2021). These low rates relative to the mesozooplankton biomass were partially due to the imminent seasonal diapause of a major fraction of the surface mesozooplankton biomass, *Neocalanus* copepods (Stamieszkin et al., 2021). Our results suggest that this decline in biomass may be in part mediated by the inability of a portion of the mesozooplankton community to meet metabolic demands with grazing in the fall, emphasizing the longer ecological time scales for this component of the ecosystem. The food web represented here exemplifies the low range of ecosystem states typical for an HNLC region.

### 4.2 Connectivity among food web processes

Correlations among daily measurements of food web components provides a mechanistic understanding of how fluctuations in primary production, which are more readily detected through remote sensing (e.g., Longhurst et al. 1995; Taboada et al. 2019), propagate through the food web. Hierarchical clustering analysis (Figure 6) suggested that changes in net primary production and phytoplankton biomass were the strongest drivers of POC variation, despite phytoplankton being a small portion (∼11%) of total POC. The variability in NPP was also closely associated with nPP, primary production driven by nitrate uptake. Changes in the rate of nPP (∼28% of NPP) were primarily responsible for driving the variability in NPP despite high regenerative production (Meyer et al., 2022). The clustering of bacterial production with nPP suggest that increases in nPP led to the production of bioavailable DOC which was followed by increased bacterial activity and bacterial biomass. Detailed analysis of bacterioplankton dynamics found that substrate availability predominantly influenced bacteria production and biomass (Stephens et al., 2020; Stephens et al., in press).

Despite the DOC pool being sufficiently large, the tight coupling between DOM production and consumption processes in the mixed layer led to a relatively small accumulation of bioavailable DOC (<5% of bulk DOC pool). Measurement uncertainties of ∼± 1 uM DOC make it difficult to relate small changes in a relatively large mixed layer bulk DOC pool to other field measurements and keep DOC from clustering with other food web variables. Independent DOM production or remineralization experiments required to assess the magnitude of DOM bioavailability were conducted on the cruise but their number was limited (Stephens et al. 2020). Thus, the fluxes into and out of the relatively large and unvarying DOC pool were largely cryptic over the time scale of this cruise (Moran et al., 2022).

Two of the loss processes of the food web, microbial respiration and microzooplankton grazing, clustered separately from the production parameters suggesting that their fluctuations are temporally disconnected from most of the other food web parameters or that high noise levels in their determination masked any causal linkages. The clustering of grazing and respiration suggests that microzooplankton grazing had the strongest influence on total heterotrophic respiration. Though temporally mismatched on daily scales, the overall balance between phytoplankton production and losses due to respiration and grazing (McNair et al., 2021) and the positive correlation between microzooplankton grazing and NCP suggests the carbon produced from fluctuations in NPP was being assimilated into the food web and contributed to heterotrophic metabolism.

Due to mismatches in temporal sample alignment and frequency (see methods for parameter selections), the correlation analysis did not include parameters that may have been important drivers of production and export. Dissolved iron (dFe) concentrations were not part of the correlation analysis but it is well-known that small changes in dFe concentrations can substantially alter rates of primary production in HNLC regions (e.g., Young et al., 1991; Harrison et al., 1999) and thus could have influenced NPP in this food web. Additionally, mesozooplankton and salps were not included but they strongly influenced the form and quantity of carbon exported from surface waters and it was the contribution of salp and euphausiid fecal pellets that most significantly altered the rate of particle export from the euphotic zone (Durkin et al., 2021; Estapa et al., 2021; Stamieszkin et al., 2021; Steinberg et al., 2022).

### 4.3 Food web structure and carbon export

The mixed layer food web was highly regenerative, no matter how the food web was interrogated, measuring the balance between gross production and respiration (-0.22 to 0.01 µmol C L^-1^ d^-1^) or accounting for the particulate carbon produced (0.07 µmol C L^-1^ d^-1^), we obtained estimates of slight production or slight consumption of carbon with error bars that span zero production. Direct measurements of bacterial growth efficiency, microbial respiration, and assessment of consumer trophic levels indicated that high rates of community respiration arose from multiple trophic transfers rather than from inefficient growth. Using the stable isotope analysis, we observed the complex trophic structures of the food web, with primary consumers (microzooplankton) occupying almost two trophic levels, making mesozooplankton 3^rd^ and 4^th^ order consumers. The multiple trophic levels within microzooplankton reflects a diverse diet that includes primary and secondary producers, including bacteria as well as other microzooplankton, while the multiple trophic levels within mesozooplankton suggest a diet including carnivory or detritivory on both micro- and mesozooplankton and minimal consumption of the primary producers. The numerous trophic links between phytoplankton and mesozooplankton gave rise to higher respiration losses, which were directly measured from bacteria, from the whole microbial community and from mesozooplankton, as well as inferred for phytoplankton (GCP-NCP) and microzooplankton. Through direct measurement of fecal pellet production rates, accounting of the production and loss of biomass, and inference of the production rate of detritus, we found that the carbon available for export from the mixed layer is primarily detrital with some small potential contribution of living cells.

The minimal amount of organic carbon that was not respired took the form of small particles of biomass or detritus. The small particles in the food web can in principle, contribute to export flux via a number of mechanisms including physical export by subduction or mixing, gravitational sinking of solitary cells, sinking of aggregates, fecal pellet production, and active export by vertical migration (Siegel et al., 2016, 2023; Steinberg & Landry, 2017). The relatively quiescent weather, intense vertical stratification, and weak horizontal density gradients during the study period (Siegel et al., 2021) did not promote export facilitated by physical mixing processes (e.g., Omand et al., 2015; Resplandy et al., 2019). Moreover, daily integrated in situ phytoplankton stocks closely matched phytoplankton stocks observed in incubation experiments, indicating that physical mixing did not substantially affect standing stocks and, by extension, export (McNair et al., 2021). Small cells and detritus can contribute to export when aggregated into larger particles. However, during the study period, no visible aggregates (>0.5 mm) were formed during experiments that quantified abiotic aggregation potential (Romanelli et al., in revision), suggesting that particle numbers were too low for aggregation to play an important role. Furthermore, no visible marine snow aggregates (> 0.5 mm) were detected in any of our Marine Snow Catcher deployments (20-500 m), indicating that the concentration of these particles was <1 per 100 L (Romanelli et al., in revision). This finding indicates that sinking phytoplankton aggregates contributed little to export from the food web.

It is the byproducts of trophic transfers, fecal pellets and detritus, that contributed to export in this food web, where the production and loss of biomass were well balanced.

Microzooplankton fecal pellets, which attenuate rapidly with depth, were a small portion (3.7%) of sediment trap particle flux as was sinking detritus (10.6% of particle flux), which appeared to be fragments of larger fecal pellets (Figure 7) (Durkin et al., 2021). The majority of the observed export flux was composed of fecal pellets from mesozooplankton (58%) and salps (27%), which each efficiently repackaged small POC into sinking particles (Figure 7) (Durkin et al., 2021; Stamieszkin et al., 2021; Steinberg et al., 2022). Salps were episodically important for export and salp fecal pellets, due to their large size and high sinking velocity, accounted for up to 72% of POC produced and exported by the whole zooplankton community in the upper 100 m at night (Durkin et al., 2021; Steinberg et al., 2022). Thus, while carbon flow within the ML was dominated by the microbial loop, these processes had only a minor contribution to carbon export. In contrast, the more variable production and loss terms associated with patchy mesozooplankton and salps had little influence on the dominant flow of carbon within the food web, but the net effect of mesozooplankton and salp grazing dominated export flux (Durkin et al., 2021; Maas et al., 2021; Stamieszkin et al., 2021; Steinberg et al., 2022). Mesozooplankton and salps which have patchy distributions in time and space (Steinberg et al., 2022) were the major outlet for carbon to leave the microbial loop.

**Figure 7:**
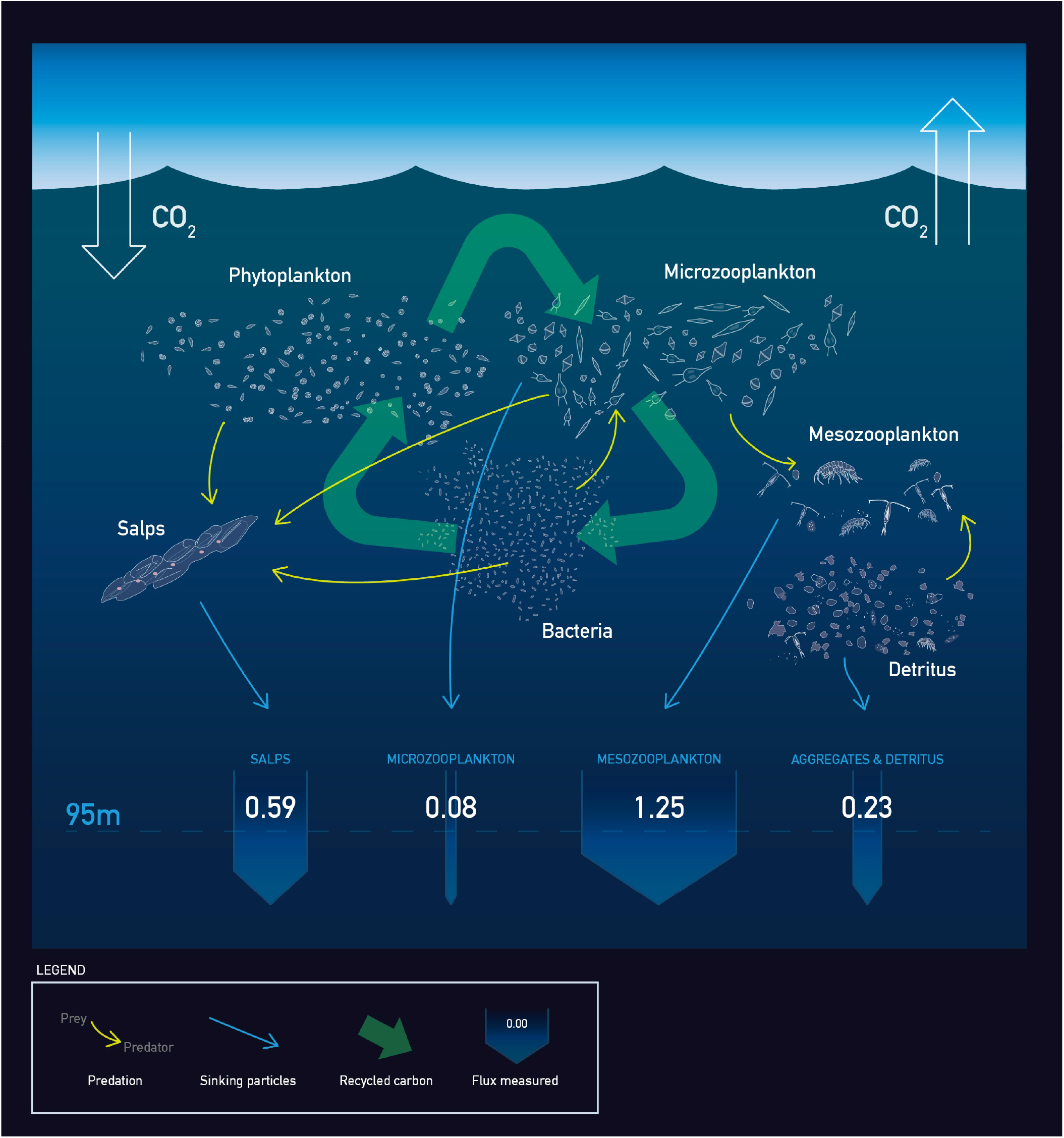
Food web graphical synopsis. The mixed layer food web was highly retentive and regenerative with the bulk of organic carbon cycled through the microbial loop and respired. Carbon removal from the microbial loop occurred via the production of fecal pellets from grazing and predation (small blue arrows). By the base of the euphotic zone (dashed line) the majority of the flux (open arrows in units of mmol C m^-2^ d^-1^) was composed of salp and mesozooplankton fecal pellets with minor contributions from microzooplankton fecal pellets and sinking detritus that appeared to be pieces of larger fecal pellets. Graphic designed by Liam Van Vleet.

## 5. Conclusion

The late summer 2018 EXPORTS campaign supported prior knowledge of the subarctic North Pacific ecosystem as a highly recycled, regenerative mixed-layer food web with low export (e.g., Bif & Hansell, 2019; Fassbender et al., 2016; Paul J. Harrison, 2002; Kirchman et al., 1993). Furthermore, by simultaneously collecting empirical data on food web stocks and rates and characterizing the quantity and quality of exported material, the EXPORTS campaign reconstructed the major pathways of carbon through the food web and out of the mixed layer, constraining uncertainties and providing turnover times. An important contrast emerged between the processes that dominated the transfer of carbon within the food web to those that contributed to export. Primary produced carbon was principally assimilated into the food web and then respired via multiple trophic transfers and microbial remineralization. In contrast, less abundant mesozooplankton that had relatively minor organic carbon uptake rates constituted the majority of the export production due to efficient repackaging of consumed material (Durkin et al., 2021; Stamieszkin et al., 2021; Steinberg et al., 2022). These low export systems are typical for much of the world’s ocean and thus suggest a comprehensive approach of measuring disparate processes is needed to quantify important carbon cycle unknowns such as the connectivity from the surface to the deep ocean organic matter reservoirs (Nowicki et al., 2022; Siegel et al., 2023). In particular, our results point to the challenging but critical need to simultaneously study microbial food web dynamics on the scales of liters and days and the processing of this microbial biomass by larger organisms within a dynamic ocean habitat spanning kilometers and weeks.

## Contributions

- Substantial contributions to conception and design: HM, MM, SL, AEM, BS, JF, TAR, MAB, DAS
- Acquisition of data: All
- Analysis and interpretation of data: All
- Drafting the article or revising it critically for important intellectual content: HM, MM, SL, AEM, BS, JF, TAR, MAB, DAS
- Final approval of the version to be published: All

## Acknowledgements

This work was made possible through the tireless efforts of the NASA EXPORTS logistics and planning teams ESPO and especially Q. Allison. The NASA program managers (P. Bontempi and L. Lorenzoni) and data management team (I. Soto Ramos and S. Craig) were responsible for making the EXPORTS project a reality. We thank the captain and crews of the R/V *Roger Revelle* and R/V *Sally Ride* for their willingness and skill dedicated to coordinating this large field campaign. For help aboard the ships, we thank S. Caprara, T. Mellett, F. Morison, and B. Ver Wey. We appreciate the artistic eye and skills of L. Van Vleet who created our summary graphic (Figure 7). This work reflects the careful and thoughtful analysis of many scientists involved in the EXPORTS project. We thank them all for sharing their data and providing helpful thoughts and suggestions that contributed to the writing of this manuscript.

## Funding Information

This work was made possible from funding from the National Aeronautics and Space Administration (NASA) and the National Science Foundation (NSF). Grant numbers and recipients are as follows:

SMD, TR: NASA 80NSSC17K0716.

SMD: NSF OCE-1736635.

AM, SG: NASA 80NSSC17K0552.

BJ, KB, MB: NSF OCE-1756442.

DS, AM: NASA 80NS SC17K0654.

CC: NASA 80NSSC18K0437.

JG: NASA 80NSSC17K0568

AS: NASA 80NSSC18K1431.

HC: NSF 1830016.

BP, KDS: NSF 1829425.

DAS, NN, KB, EF, SK, AB, AM, UP: NASA 80NSSC17K0692.

KB, CBN, LR, MRM: NASA 80NSSC17K0555.

ME, KB, CD, MO: NASA 80NSSC17K0662.

---

**Supplemental Figure 1:** Pearson’s correlation matrixes. Color coded Pearson’s correlation coefficients (rho), red indicates positive correlation and blue squares indicates negative correlation. Significant correlations (p-value <0.05) are noted with labeled R values. Correlations among two data points were replaced with gray NA values. (a) Correlations for all available food web data at daily resolution (b) Correlations for the subset of data used to create (Figure 6). Abbreviations are as follows: net primary production (NPP), chlorophyll a (chl *a*), new primary production (nPP), net community production (NCP), microzooplankton grazing rate (µzoop grazing), integrated daily radiation (I_g_), ortho phosphate (PO_4_^3-^), microbial community respiration (CR), particulate organic carbon (POC), nitrate (NO_3_^-^), silicate (Si(OH)_4_), and dissolved organic carbon (DOC).

