## Supplemental Figure 1 for "Quantitative analysis of food web dynamics in a low export ecosystem"

a

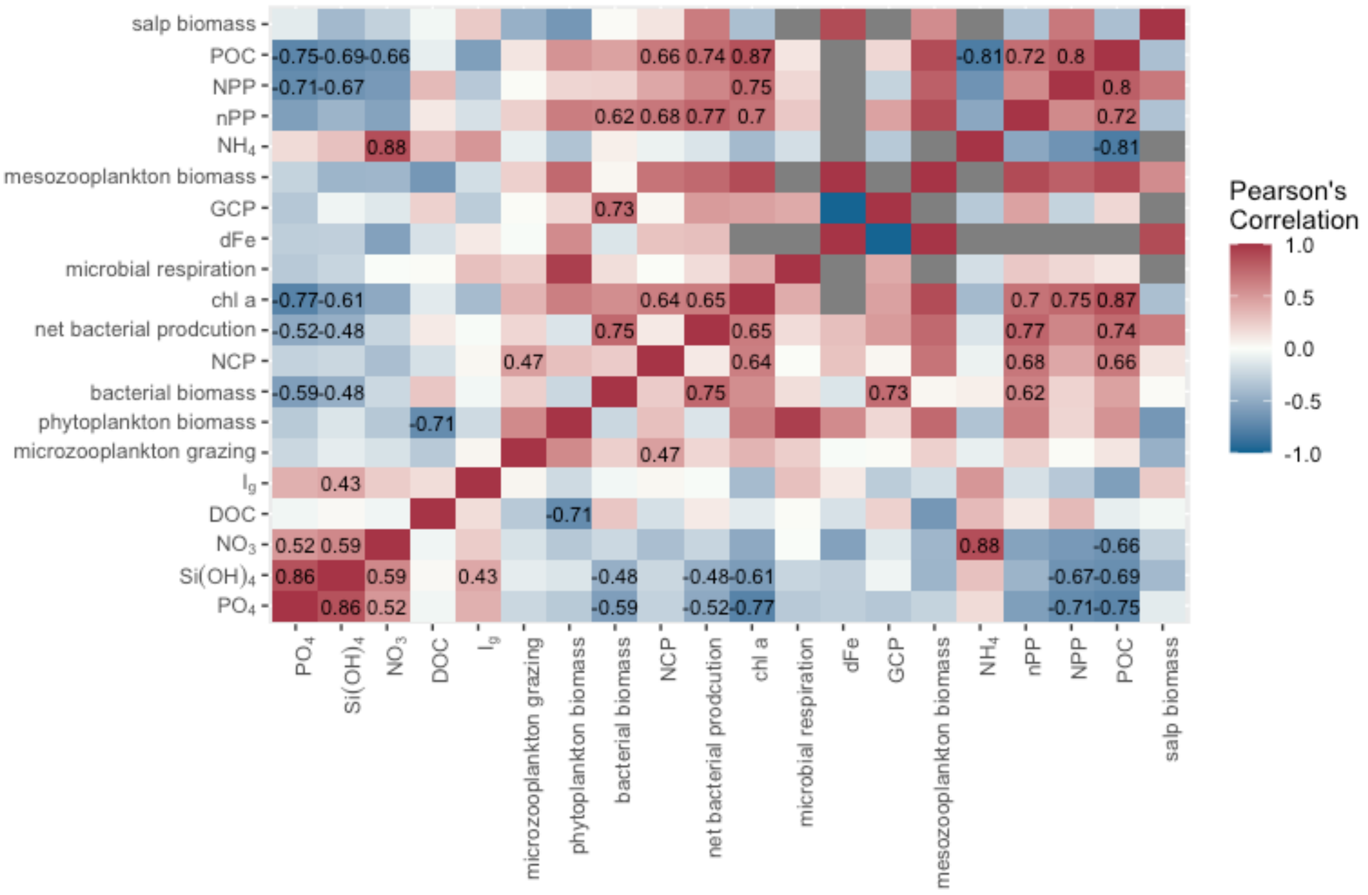


b

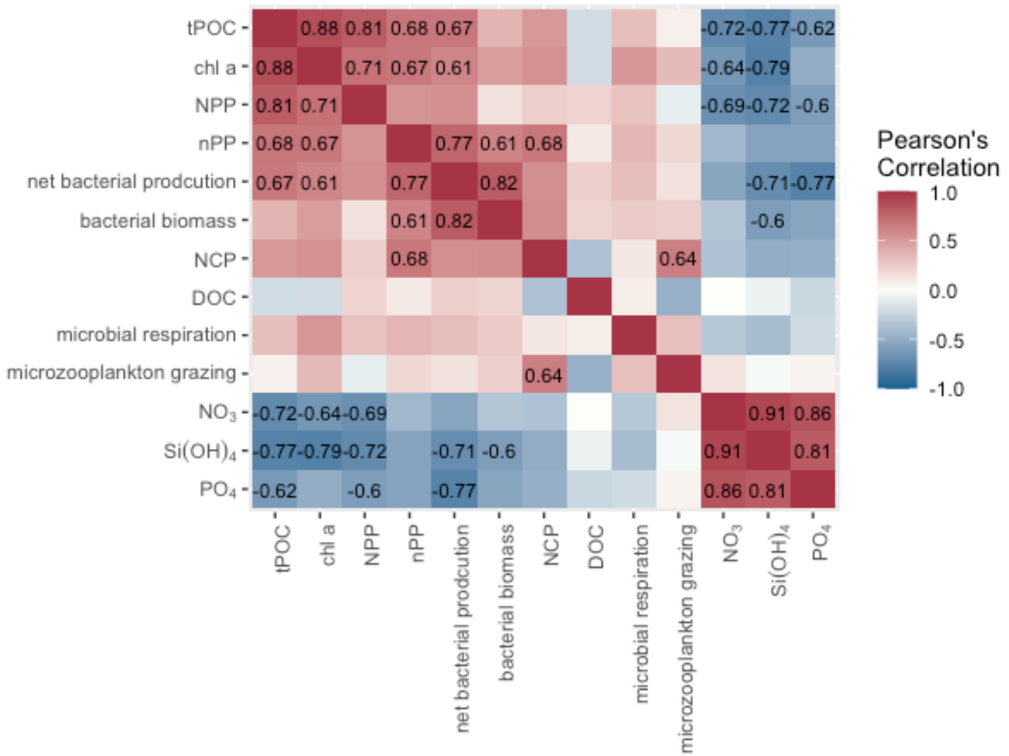

**Supplemental material**

**Dissolved Fe measurements**

Surface (~2 m) water samples for dissolved iron (Fe) concentrations were collected using a custom trace metal clean “towfish” sampling system (Mellett and Buck 2020) on the *R/V* Roger Revelle between 18 August and 6 September 2018. A total of 24 samples were filtered (<0.2 µm, Pall Acropak) inline, collected in acid-cleaned 125-mL low density polyethylene (LDPE, Nalgene) bottles, and acidified to 0.024 M hydrochloric acid (HCl, Fisher Optima). Dissolved trace metal Fe concentrations were determined by high resolution inductively coupled plasma mass spectrometry (HR-ICP-MS) at the University of South Florida (Hollister et al. 2020). The limit of detection (LOD) of this method was 0.028 nM for dissolved Fe.

**Microzooplankton and Phytoplankton community genetic composition**

Grazer community structure was determined by with high-throughput sequencing of the small ribosomal subunit 18S rDNA gene of eukaryotes. Total DNA was extracted from plankton biomass samples using DNeasy Blood and Tissue kit (Qiagen). DNA concentration was quantified with NanoDrop Spectrophotometer (ThermoFisher). The V4 hypervariable region of 18S rDNA gene was amplified using primers Reuk454FWD1 and V4r which amplify all eukaryotes (Bradley et al., 2016). Primers were modified by the addition of Illumina specific adapters: Reuk_454FWD1 5'-TCG TCG GCA TCA GAT GTG TAT AAG AGA CAG CCA GCA SCY GCG GTA ATT CC-3 and V4r 5'-GTC TCG TGG GCT CGG AGA TGT GTA TAA GAG ACA GAC TTT CGT TCT TGA T-3'. The following reagents were used in 35 µL PCR reactions; Accustart ll PCR mix 2X (VWR 89235-018), 10 µM each forward and reverse primer, and (include approx. ng) 3uL DNA template. Reactions were amplified with a multi-step thermocycler protocol, consisting of a 2 min denaturing step at 94 °C, followed by 30 cycles of 30 s each at 94, and 55, 1 minute 72 °C, followed by 10 minutes at 72 °C.

PCR amplicons were cleaned with Ampure XP beads (Beckman Coulter, Inc., Brea, CA, USA), quantified with the Qubit High Sensitivity DNA Assay Kit (Thermo Fisher Scientific, Inc., Waltham, MA, USA), amplified for an additional five cycles to add Nextera indices and adaptors (Illumina, Inc., San Diego, CA, USA) and cleaned again with Ampure XP beads. PCR products were pooled and quantified with the KAPA qPCR kit (Kapa Biosystems, Wilmington, MA, USA) prior to Illumina MiSeq sequencing with V3 chemistry (2 × 300 bp reads; Illumina, Inc., San Diego, CA, USA) at the University of Rhode Island Genomics and Sequencing Center.

**Bioinformatic pipeline and downstream analysis**

Bioinformatics of the 18S rRNA high-throughput sequences were conducted using the DADA2 package (Callahan et al. 2016; version 1.16) and visualized with Phyloseq (McMurdie and Holmes 2013). Paired end sequencing reads were first trimmed to remove low quality bases and Illumina adaptors using the CUTADAPT package (Martin, 2011), followed by a 4 bp sliding window quality score of 20 and a minimum read length of 200 bp. Trimmed sequences were filtered to a minimum length of 225 and 210 bp for forward and reverse reads, respectively; and a maximum expected error (based on phred quality scores) of 2. Paired end sequences were merged using the mergePairs function from DADA2. After appropriate quality control, merged sequences were assigned taxonomy using the assignTaxonomy function of DADA2 and the PR2 Database (pr2_version_4.12.0). The resulting amplicon sequence variants (ASVs) were processed to remove ASVs that contained fewer than 0.4% total sequence reads per sample to control for sequencing error and to obtain meaningful ecological information.

$k = \mu - g$

We solve for g by taking the difference of the net growth rate in the diluted and undiluted treatments and dividing by the dilution factor, in the case of a 30% sea water to 70% filtered seawater dilution. Rearranging we then get

$g = (k_{diluted} - k) / (1-0.3)$

**Trophic position using stable isotopes**

Collection of zooplankton for CSIA-AA:

The contents of the cod end were split and one quarter was separated by size fraction using nested sieves of 5.0, 2.0, 1.0, 0.5, and 0.2 mm mesh, and then rinsed with filtered seawater follower by isotonic ammonium formate over pre-weighed 0.2 mm mesh nitex filters before being frozen at -20° C for biomass and stable isotope analysis.

Analysis:

Zooplankton samples were prepared for amino acid stable isotope analysis using routine using routine procedures outlined in Hannides et al. (2009). The nitrogen isotopic composition of zooplankton was measured using a Thermo Scientific Delta V Plus IRMS interfaced to a Trace gas chromatograph fitted with a 60 m BPx5 capillary column (SGE Analytical Science, 0.32 mm diameter, 1 µm stationary phase) via a GC-C III combustion interface (980 °C) with reduction furnace (650 °C)) and liquid nitrogen cold trap. Internal reference compounds, L-2-Aminoadipic acid (AAA) and L-(+)-Norleucine (NOR) of known nitrogen isotopic composition, were co-injected with samples and used to determine accuracy and precision. During analysis of zooplankton, a suite of 14 pure amino acids of known isotopic composition were co-injected with noroleucine and aminoadipic acid and measured every 2-4 sample injections as a reference material. The results from analyses of the amino acids suite were used to derive a linear correction to normalize measured sample amino acids $\delta^{15}$N values. Corrected results are reported in δ-notation relative to atmospheric $N_2$. Precision of determination of $\delta^{15}$N values was ±0.3‰.

Trophic position was calculated from $\delta^{15}\mathrm{N}_{\mathrm{AA}}$ values using the equation:

$$TP = \frac{\delta^{15}\mathrm{N}_{tr} - \delta^{15}\mathrm{N}_{src} + \beta_{tr-src}}{TDF_{tr-src}} + 1$$

where $\delta^{15}\mathrm{N}_{\mathrm{tr}}$ and $\delta^{15}\mathrm{N}_{\mathrm{src}}$ are the $\delta^{15}$N values associated with some specified individual or set of trophic and source amino acids, $\beta$ is the difference between $\delta^{15}\mathrm{N}_{\mathrm{tr}}$ and $\delta^{15}\mathrm{N}_{\mathrm{src}}$ in primary producers, and $TDF$ (the trophic discrimination factor) describes how much $\delta^{15}\mathrm{N}_{\mathrm{tr}}$ changes relative to $\delta^{15}\mathrm{N}_{\mathrm{src}}$ with each trophic step. Both $\beta$ and $TDF$ were determined empirically, and specific to the set of amino acids used. $TP$ was estimated using two different sets of amino acids, which are tabulated along with their respective $\beta$ values and $TDF$s relative to phenylalanine in the below table. $TP_{ala-phe}$ was used to estimate total food web length, inclusive of protistan heterotrophy, as in Décima and Landry (2020). Because error was not reported in Décima and Landry (2020), estimates of error in $TDF_{ala-phe}$ and $\beta_{ala-phe}$ from Décima et al. (2017) were used and are considered conservative. $TP_{glu-phe}$ was used to estimate the length of the metazoan food web, exclusive of protistan heterotrophy, as in Chikaraishi et al. (2009). These two estimates of $TP$ were then compared by subtracting $TP_{glu-phe}$ from $TP_{ala-phe}$ to obtain $\Delta TP_{ala-glu}$, in order to estimate the length of the protistan food web, and error was propagated through that calculation. $TP_{tr-src}$ is also reported here and was calculated as in (Jarman et al. 2017) and (Ohkouchi et al. 2017).

| | TDF | beta |
|---|---|---|
| Glx | 7.1 $\pm$ 1.2 ‰ | 3.4 $\pm$ 0.9 ‰ |
| Ala | 4.5 $\pm$ 2.1 ‰ | 3.2 $\pm$ 1.2 ‰ |
